# DNA methylation carries signatures of sublethal effects under thermal stress in loggerhead sea turtles

**DOI:** 10.1101/2023.11.22.568239

**Authors:** Eugenie C. Yen, James D. Gilbert, Alice Balard, Inês O. Afonso, Kirsten Fairweather, Débora Newlands, Artur Lopes, Sandra M. Correia, Albert Taxonera, Stephen J. Rossiter, José M. Martín-Durán, Christophe Eizaguirre

**Author notes:** **Corresponding Author:** Eugenie C. Yen. School of Biological and Behavioural Sciences, Queen Mary University of London, London, E1 4DQ, UK.

## Abstract

Rising global temperatures are a major threat to biodiversity. Whilst research generally focuses on thermal tolerance and mortality, sublethal effects may alter population dynamics and subsequently the adaptive potential of species. However, detecting such effects in the wild can be challenging, particularly for endangered and long-lived species with cryptic life histories. This necessitates the development of molecular tools to identify their signatures. In a split-clutch design experiment, we relocated clutches of wild, nesting loggerhead sea turtles (*Caretta caretta*) to a protected, *in-situ* hatchery. Eggs were then split into two sub-clutches incubated under shallow (35cm) or deep (55cm) conditions, with those in the shallow treatment experiencing significantly higher temperatures. Whilst no difference in hatching success was observed between treatments, hatchlings that emerged from the shallow, warmer treatment had altered length-mass relationships, and were weaker at fitness tests of locomotion capacity than their siblings incubated in the deep, cooler treatment. To characterise the molecular signatures of these thermal effects, we performed whole genome bisulfite sequencing on blood samples collected upon emergence. This identified 714 differentially methylated sites between treatments, including on genes with neuronal development, cytoskeleton, and sex determination functions. Taken together, our results show that higher incubation temperatures can induce sublethal effects in hatchlings, which are reflected in their DNA methylation status at identified sites. Such sites could be used as biomarkers of thermal stress, especially if they are retained across life stages. Overall, this study suggests that global warming may have population-level consequences for loggerhead sea turtles, by reducing hatchling quality, dispersal capacity and the adaptive potential of this species. Conservation efforts for climate-threatened taxa like endangered sea turtles will therefore benefit from strategies that monitor and mitigate exposure to incubation temperatures that lead to sublethal effects.

## Introduction

The current pace of biodiversity loss is often referred to as the sixth mass extinction event in geological history (Barnosky et al., 2011). As global temperatures rise rapidly, conservation programmes that promote the persistence of climate-vulnerable taxa are urgently needed (Stocker et al., 2013; Urban 2015). To be effective, these mitigation strategies must integrate knowledge on how target species interact with their environment, to accurately assess extinction risk and guide appropriate interventions within their unique, eco-evolutionary contexts (Lavergne et al. 2010; Hoffmann & Sgrò, 2011; Urban et al. 2016).

Species that avoid extinction typically follow two pathways (Aitken et al 2008). They can shift in distribution (Chen et al. 2011; Lenoir & Svenning, 2015), usually towards the poles (Antão et al 2022), higher elevation (Mamantov et al. 2021) or greater depth in aquatic systems (Pinsky et al. 2020). Alternatively, species may adapt *in-situ* via genetic evolution if effective population size and reproduction rates are sufficiently high, through adaptive plastic responses, or a context-dependent combination of both (Franks & Hoffmann, 2012; Catullo et al. 2019). Despite those available mechanisms, responses may still be too slow to keep up with the current rate of climate change for some species (Radchuck et al 2019). Both the ecological range-shift and evolutionary adaptive scenarios focus on the longer-term response to temperature increase. Yet, an immediate threat relates to the impact of sublethal physiological effects on individuals (Pörtner et al., 2006; Carlo et al 2018; Conradie et al 2019). Characterising these effects is important in a conservation context, as negative fitness-impacts on ecologically relevant functions can exacerbate extinction risk by altering population dynamics, such as reproduction and recruitment rates (Hamilton et al. 2016). Their characterisation and integration will therefore strengthen predictive models and enhance the development of targeted mitigation strategies for threatened species (Chown et al. 2010; Lockley and Eizaguirre, 2020).

Sublethal effects however can be difficult to measure in the wild, particularly in animal species with complex and cryptic life history stages. To overcome this challenge, conservation can benefit from molecular approaches employed in medicine, where biomarkers are frequently developed to understand and diagnose sublethal effects in patients (Silins and Högberg, 2011; Sarhadi and Armengol, 2022). Biomarkers are also common in ecotoxicology for characterising sublethal exposure to pesticides and pollutants (Sarkar et al. 2006; Forbes et al. 2006; Vischetti et al. 2020). To this end, we need to identify relevant molecular signatures associated with thermal, sublethal effects in threatened species. Whilst proteomic profiles can be altered by exposure to elevated temperatures (Abdelnour et al. 2019), application is hindered by labour-intensive protocols. Gene expression levels are often associated with heat stress, however unstable RNA can degrade rapidly post-sampling, rendering it a complex molecule to implement as a field diagnostic marker (Desalvo et al. 2008; Lim et al. 2016; Akbarzadeh et al. 2018). An alternative is to harness the potential of DNA methylation, which is an epigenetic regulatory mechanism involving the addition of a methyl group to cytosine residues (Rey et al. 2020).

Quantifying DNA methylation is a feasible option in the field as it can stably maintain regulatory information (Gosselt et al. 2021). Induction of methylation changes following exposure to environmental stressors has been widely shown (Metzger and Schulte, 2017; Guan et al. 2019; Sheldon et al. 2020), with links to growth, metabolism, and behaviour (Guerrero-Bosagna et al. 2020; Sepers et al. 2021). DNA methylation can also integrate environmental effects across life stages (Pértille et al. 2017; Jonsson and Jonsson, 2019; Bock et al. 2022), and even across generations in some cases (Blaze and Roth, 2015; Heckwolf et al. 2020). Taking these attributes together, DNA methylation profiling presents a promising approach to identify molecular biomarkers that link thermal stress to variation in functional traits (Jeremias et al. 2020; Šrut, 2021; Crossman et al. 2021), thus providing a practical solution to monitoring sublethal effects on wild populations (Rey et al. 2020).

Sea turtles are ectothermic species facing risks from warming oceans and nesting beaches due to their life cycle across both environments (Wallace et al. 2011). Vulnerability is further heightened by their temperature-dependent sex determination system (Janzen, 1994), which may feminise sex ratios to the point of demographic collapse under global temperature projections (Hawkes et al. 2009; Mitchell & Janzen, 2010). Whilst much research focuses on the impact of warming on sex ratios, increasing incubation temperatures can also impose fitness costs on individuals, which could weaken the adaptive potential of the population and accelerate their decline (Eizaguirre and Baltazar-Soares, 2020). For example, even though incubation temperatures above 34°C are often considered to be deadly for sea turtle nests (Howard et al. 2014), hatchlings that emerge from nests incubated at low temperatures (27°C) or above 33°C experience slower growth rates than those incubated at around 30°C (Booth et al. 2020). High temperatures can also weaken crawling, self-righting and swimming ability, thereby reducing dispersal capacity toward the oceans after hatching (e.g. Booth and Evans, 2011; Fisher et al. 2014; Mueller et al. 2019).

In this study, we focus on the endangered loggerhead sea turtles (*Caretta caretta)* that nest in Cabo Verde, West Africa. This is one of the most threatened populations for this species due to coastal development, sea level rise and poaching (Taxonera et al. 2022). The population is composed of distinct genetic groups, arising from strong female philopatry down to just tens of kilometres (Stiebens et al. 2013; Baltazar-Soares et al. 2020). Current debate exists regarding the future of the Cabo Verde population in relation to rising incubation temperatures, with modelling studies predicting almost complete feminisation by the century end (Laloë et al. 2014), alongside other studies arguing that this population will be resilient (Abella Perez et al. 2016). This debate, however, does not consider how sublethal, fitness costs on individuals will impact the resilience potential of this population, nor whether there are biomarkers that can be used to monitor such effects.

To begin addressing this challenge, we conducted a split-clutch design experiment in an *in-situ* hatchery, exposing clutches of wild, nesting females to different incubation temperatures under field-relevant conditions. This was achieved by burying each sub-clutch at either 55cm (deep treatment) or 35cm (shallow treatment), which exposed shallow sub-clutches to increased temperatures that mimic future climate projections in Cabo Verde. Using a split-clutch design enabled us to characterise developmental stress responses associated with incubation conditions, independently of the maternal genetic background. Upon emergence, a small blood sample was collected for whole genome bisulfite sequencing to identify differentially methylated sites between treatments that could be used as biomarkers of incubation condition. Fitness tests related to hatchling quality and locomotion capacity were also conducted and linked to methylation values at genomic sites of interest. Within the context of regional climate projections, we consider the implications of our results for the persistence of this genetically distinct and globally important loggerhead sea turtle population.

## Material and Methods

### 1. Experimental design

Our study focused on loggerhead sea turtles that nest on Sal Island, which is the most north-easterly island of the Cabo Verde Archipelago, in the Northeast Atlantic. There, the sea turtle nesting season runs from late June to October. The sampling site of Algodoeiro Beach (16.61773 °N, −22.92882 °E) covers 800m of sandy coastline on the lower southwest of the island. On the 29^th^ of July 2021, we relocated the clutches of ten wild, nesting females over one night, to standardise environmental conditions for all egg clutches during incubation. Immediately after oviposition, females were individually marked with a passive integrated transponder (PIT) tag on their front right flipper for identification (Stiebens et al. 2013). At the end of the natural nesting process, clutches of these females (82 ± 16 (standard deviation, SD) eggs) were then relocated to an experimental *in-situ* hatchery protected from predation.

In the hatchery, we set up a split-clutch design experiment to understand the sublethal effects of incubation temperature in a field setting, whilst controlling for genetic background and maternal effects (Eizaguirre et al. 2012). Each clutch was randomly split into two sub-clutches of equal size, hence also controlling for metabolic heating, then buried at different depths to generate different incubation temperatures. One sub-clutch was buried at 55cm as the ‘deep’ treatment, which is lower than the maximum nest depth in the original nesting site. The other sub-clutch was buried at 35cm as the ‘shallow’ treatment, thereby raising the incubation temperature to mimic future conditions predicted for this population (Laloë et al. 2014). A HOBO Pendant MX Water Temp™ (MX2201 model) temperature logger was placed at the centre of 16 out of 20 sub-clutches and programmed to take a reading every 15 minutes throughout the incubation period (accuracy ± 0.5°C). Total incubation duration was calculated in days from the date each nest was closed, to the date where most hatchlings emerged from the nest simultaneously. Following natural emergence, nests were excavated and the number of unhatched or dead hatchlings were counted, as well as any remaining live hatchlings that were released onto the beach for natural dispersal. Hatching success was calculated as the ratio of live hatchlings over the total number of eggs per sub-clutch.

### 2. Hatchling morphometrics and locomotion tests

Upon emergence, up to twenty hatchlings per sub-clutch (n=408 in total, n=198 from deep treatment, n=210 from shallow treatment) were randomly selected and measured for two morphometric traits. These were the: (1) notch-to-notch straight carapace length (SCL, in millimetres, mm), and (2) mass (in grams, g). SCL was recorded as the mean of three measurements per hatchling using a digital calliper (± 0.01 mm), ensuring all measurements fell within a range of 0.5 mm. Mass was measured once per hatchling using a digital scale (± 0.1 g).

In addition, two fitness tests of locomotion capacity were conducted: (1) run time and (2) self-righting time. These traits are important components of predator and obstacle avoidance, which are both required to successfully reach the ocean after emerging from the nests (Scott et al 2014, Lockley et al 2020; Martins et al. 2020). We measured run time as the time taken for a hatchling to crawl along a 0.5 m runway of flat sand between two wooden pieces, with a dull red light at the end of it. This trial was repeated twice per hatchling. If a hatchling did not attempt to crawl at all, it was considered to have failed the trial. The mean run time (in seconds, s) was then calculated across the successful trials per hatchling. Self-righting time was measured by placing a hatchling on its back on an area of flat sand, and timing how long it took to right itself. This trial was repeated three times per hatchling. If a hatchling took longer than 1 min to self-right, it was considered to have failed the trial. We then measured the mean self-righting time (s) using the successful trials per hatchling.

### 3. Clutch- and hatchling-level phenotype analyses

All statistical analyses on phenotypic traits were carried out in RStudio v.4.1.1 (R Core Team, 2021). These were conducted at two levels. Firstly, at the sub-clutch level, we tested whether depth treatment was associated to different mean incubation temperatures with a linear model. Using the same test, we also investigated whether depth treatment was associated with different incubation durations. We then tested if hatching success rates correlated with full clutch size (and its quadratic term), mean sub-clutch incubation temperature, depth treatment, as well as all two-ways interactions with the depth treatment variable. To account for the correlation between depth treatment and temperature, we calculated the residuals of a linear model between those two variables and used them in the main model to replace the mean sub-clutch incubation temperature.

Secondly, we focused on the individual-level data for hatchlings. We used a series of linear mixed-effects models with maternal ID included as a random effect, in the R package lme4 v.1.1-33 (Bates et al. 2014). Specifically, we investigated correlates of hatchling size (SCL), considering their mass, depth treatment, clutch size and mean sub-clutch incubation temperature. Models were stepwise selected using the step function of the R package lmerTest (Kuznetsova et al. 2017). Mean run and self-righting times were log_10_+1 transformed and tested for correlations with clutch size, depth treatment and mean sub-clutch incubation temperature, as well as their interactions. Hatchling SCL was used as a covariable, and maternal ID was set as a random factor. All figures were made using the R package ggplot2 v.3.4.2 (Wickham, 2016).

### 4. Sample collection for molecular work

Immediately after morphometric measurements and completing fitness tests, 100 μl of blood was sampled from the dorsal cervical sinus of each hatchling, using a 26-gauge needle and 1 ml syringe (Wibbels et al. 1998). Collected blood samples were stored in lithium heparin coated tubes and refrigerated for up to 12 hours. Samples were then centrifuged for 1 min at 3000 rpm to separate plasma and red blood cells. These samples were stored at −18°C until the end of the field season, then at −80°C following transport to Queen Mary University of London (London, UK).

### 5. DNA extraction and sequencing

Red blood cell samples from a subset of 40 hatchlings (20 hatchlings per deep and shallow treatment) were selected for whole genome bisulfite sequencing (WGBS), with two hatchlings sequenced per sub-clutch. Within each sub-clutch, a hatchling was chosen for sequencing if it had not failed any fitness tests, with priority given to hatchlings that were sampled earlier in the sub-clutch, to reduce effects of waiting time on fitness test performance. This approach is as conservative as it can be regarding sublethal effects, since it removes the weakest hatchlings. Genomic DNA was extracted from the 40 red blood cell samples using a QIAGEN Blood and Tissue Kit (Qiagen, Germany), following the manufacturer’s protocol. Sodium bisulfite treatment, library preparation and WGBS was performed by BGI (Hong Kong). DNBseq whole genome bisulfite libraries were constructed and sequenced with 100 base pair (bp) paired end reads on an MGI DNBSEQ platform (BGI, Hong Kong). This generated a mean of 134,492,469 ± 3,641,276 (SD) reads per sample (**Supplementary Table 1**).

### 6. Read trimming and methylation calling

Raw WGBS reads were trimmed for residual adapters and filtered for a mean Phred score of higher than Q20 using cutadapt v.2.10 (Martin, 2011). Read quality control was conducted pre and post trimming with FastQC v.0.11.9 (Andrews, 2010). Using Bismark v.0.22.1 with default options (Krueger and Andrews, 2011), trimmed reads were aligned against our novel, chromosomal-scale reference genome assembly for a loggerhead sea turtle from Sal Island, Cabo Verde (Yen et al. in prep). This gave a mean mapping efficiency of 78.5 ± 3.4 (SD) % per sample (**Table S1**). Alignments were deduplicated with Bismark in paired end mode, followed by merging and sorting with samtools v.1.9. (Li et al. 2009). Methylation calling was then performed in Bismark. Percentage methylation totalled across CHG and CHH sites were calculated to obtain an estimate of bisulfite conversion efficiency (Laine et al. 2023). This gave a mean of 99.993 ± 0.002 (SD) %, supporting high conversion efficiency (**Table S1**). We focused on methylation patterns at CpG sites for our study, since this is the major methylation context in vertebrates (Klughammer et al. 2023). To improve coverage and minimise pseudo-replication, we further de-stranded adjacent cytosines per CpG site using the ‘merge_CpG.py’ script (Cristofari, 2023), as methylation occurs symmetrically at CpG sites in vertebrates (Klughammer et al. 2023). On average, this resulted in 24,947,580 ± 825,959 (SD) CpG sites per sample, with a de-stranded coverage of 8.56 ± 0.93 (SD) X (**Table S1**).

Methylation calls at CpG sites were further processed in RStudio v.4.2.2 (R Core Team, 2021) with the methylKit package v.1.24.0 (Akalin et al. 2012). CpG sites were excluded if they had a coverage lower than 5X, or if they were within the 99.9th percentile to account for possible PCR bias (Wreczycka et al. 2017). Coverage was then normalised between samples using methylKit’s ‘normalizeCoverage’ function. Following coverage filtering, we implemented a stringent filter that only passed CpG sites covered in all individuals (n=40), leaving 2,739,804 sites. This is because we are interested in testing whether there are sites which can be used in the future as universal biomarkers for this population. As a final filtering step, CpG sites that were potential C-to-T single nucleotide polymorphisms (SNPs) were removed, because these can be misidentified as non-methylated Cs in bisulfite-treated DNA, and subsequently bias differential methylation estimates (Wreczycka et al. 2017). To obtain a list of C-to-T SNP sites to remove, the Revelio algorithm was used to mask bases generated by the bisulfite conversion process that may be incorrectly interpreted as SNPs (Nunn et al. 2022), then the GATK pipeline v.4.2.6.1 (McKenna et al. 2010) was used to call SNPs (**Text S1** for full methods). We removed all CpG sites sharing genomic coordinates with identified C-to-T SNPs, leaving 2,733,573 CpG sites (99.77%) for downstream analyses.

### 7. Global methylation analyses

To characterise changes in the entire DNA methylome in response to incubation treatment, we compared global methylation patterns between hatchling from the deep and shallow treatments at 2,733,573 CpG sites. Using methylKit’s ‘percMethylation’ function, matrices of percentage methylation values per individual at each CpG site were generated. Methylation level per individual was obtained by counting the number of sites that exhibited (1) non-zero methylation (>0%), and (2) a methylation percentage greater than 70% (Sagonas et al. 2020). Correlations between methylation count and treatment were tested using linear mixed-effect models with maternal ID included as a random effect, in the R package lme4 (Bates et al. 2014). We further performed cluster analyses of global methylation patterns, with sites of low variation across individuals (SD<0.3) excluded, since these are non-informative for clustering. Hierarchical clustering was performed using methylKit’s ‘clusterSamples’ function, with the Ward agglomeration method and correlation distance method. Non-metric multidimensional scaling plots (NMDS) were also computed using the Bray-Curtis dissimilarity matrix with k=6 dimensions and a maximum of 1000 iterations, through the ‘metaMDS’ function of the R package vegan v.2.6.4 (Oksanen et al. 2017). To assess the contribution of treatment group and maternal ID to methylation variance, permutational multivariate analysis of variance (PERMANOVA) was conducted using vegan’s ‘adonis2’ function with 999 permutations.

### 8. Identification of DMS between incubation treatments

To identify differentially methylated sites (DMS) between individuals incubated in the deep and shallow treatments, we used the ‘calculateDiffMeth’ function of methylKit to produce differential methylation statistics. This involved performing a logistic regression test per CpG site between treatment groups with maternal ID set as a covariate, and the sliding linear model method (SLIM) for multiple testing correction (Wang et al. 2011). Methylation difference at each CpG site was calculated as the mean difference between hatchlings from the deep (control) treatment minus the shallow treatment, weighted by read coverage (Akalin et al. 2012). We considered a CpG site to be differentially methylated if it had a methylation difference greater than 15% between hatchlings from the two treatments, and a q-value less than 0.01. We further performed hierarchical clustering, NMDS and PERMANOVA analyses with the subset of identified DMS sites only, using the same parameters as described previously for global methylation analyses.

### 9. Annotation of genomic location and genes associated with DMS

To annotate the type of genomic region upon which CpG sites reside, we used the R packages genomation v.1.30.0 (Akalin et al. 2015) and GenomicRanges v.1.50.2 (Lawrence et al. 2013) alongside our reference genome annotation (Yen et al. in prep). Promoter regions were defined as 1500 bp upstream and up to 500 bp downstream of a transcriptional start site (TSS, Heckwolf et al. 2020). All CpG sites were assigned to one of four genomic categories using genomation’s ‘annotateWithGeneParts’ function, in the following order of precedence when features overlapped: promoter, exon, intron or intergenic region. A chi-squared test was used to evaluate whether DMS were differently distributed across genomic region types compared to all CpG sites.

To attach functional gene information to our DMS, those in genic regions (i.e., located on a promoter, intron or exon) were associated to a gene using the ‘findOverlaps’ function of GenomicRanges. DMS in intergenic regions were associated to a gene using genomation’s ‘getAssociationWithTSS’ function, if they were less than 10 kb away from the nearest TSS (Heckwolf et al. 2020). We further manually selected DMS of interest to investigate their gene functions deeper. These were identified based on their methylation and functional annotation characteristics. We focused on DMS identified between individuals incubated in the two treatment groups, which were: (1) located on genes with at least three DMS, (2) the top 5 most differentiated DMS, (3) located on promoters with the most DMS, and (4) the most differentiated DMS in both directions on promoters. This resulted in 21 DMS of interest (**Table S3**).

### 10. Correlating methylation value at DMS of interest against fitness-related phenotypes

To investigate the link between hatchling phenotype and molecular responses to incubation conditions, we tested for correlations between hatchling fitness-related traits and methylation values at the 21 DMS of interest. We performed a series of independent linear mixed effect models testing for the interaction between methylation value at the DMS with incubation treatment, on hatchling SCL, mass, run and self-righting times. Maternal clutch ID was set as a random factor. P-values were corrected for multiple testing (21 DMS of interest x 4 fitness traits) using the false discovery rate (FDR) method with a p-value of 0.0097.

### 11. Functional enrichment analyses

For all genes associated with DMS that had attached Gene Ontology (GO) terms in the loggerhead sea turtle genome annotation (n=247 genes, Yen et al. in prep), we performed a conditional hypergeometric GO term enrichment analysis, using the R packages GOstats v.64.0 (Falcon and Gentleman, 2007) and GSEABase v.1.60.0 (Morgan et al. 2023). The gene sub-universes were assigned as genes associated to DMS, split into hyper-methylated (i.e., higher methylation values in deep-incubated hatchlings, n=110) and hypo-methylated sites (i.e., higher methylation value in shallow-incubated hatchlings, n=137). These were compared against the gene universe, which was set as all genes associated to CpG sites present in all individuals with GO terms available (n=13,681 genes). Overrepresented biological processes, molecular functions and cellular components were then identified, using the FDR method with a threshold of 0.01 for multiple testing correction.

## Results

### 1. Clutch-level phenotypes

The incubation period lasted 53.6 days on average (minimum 49 days, maximum 58 days), over which the mean temperatures ranged from 30.19°C to 31.60°C (**Figure 1A**). Incubation duration and mean incubation temperature per sub-clutch were strongly negatively correlated (R^2^=0.88, F1,14=115.49, p<0.001, **Figure S1A**), as expected for this species.

**Figure 1.**
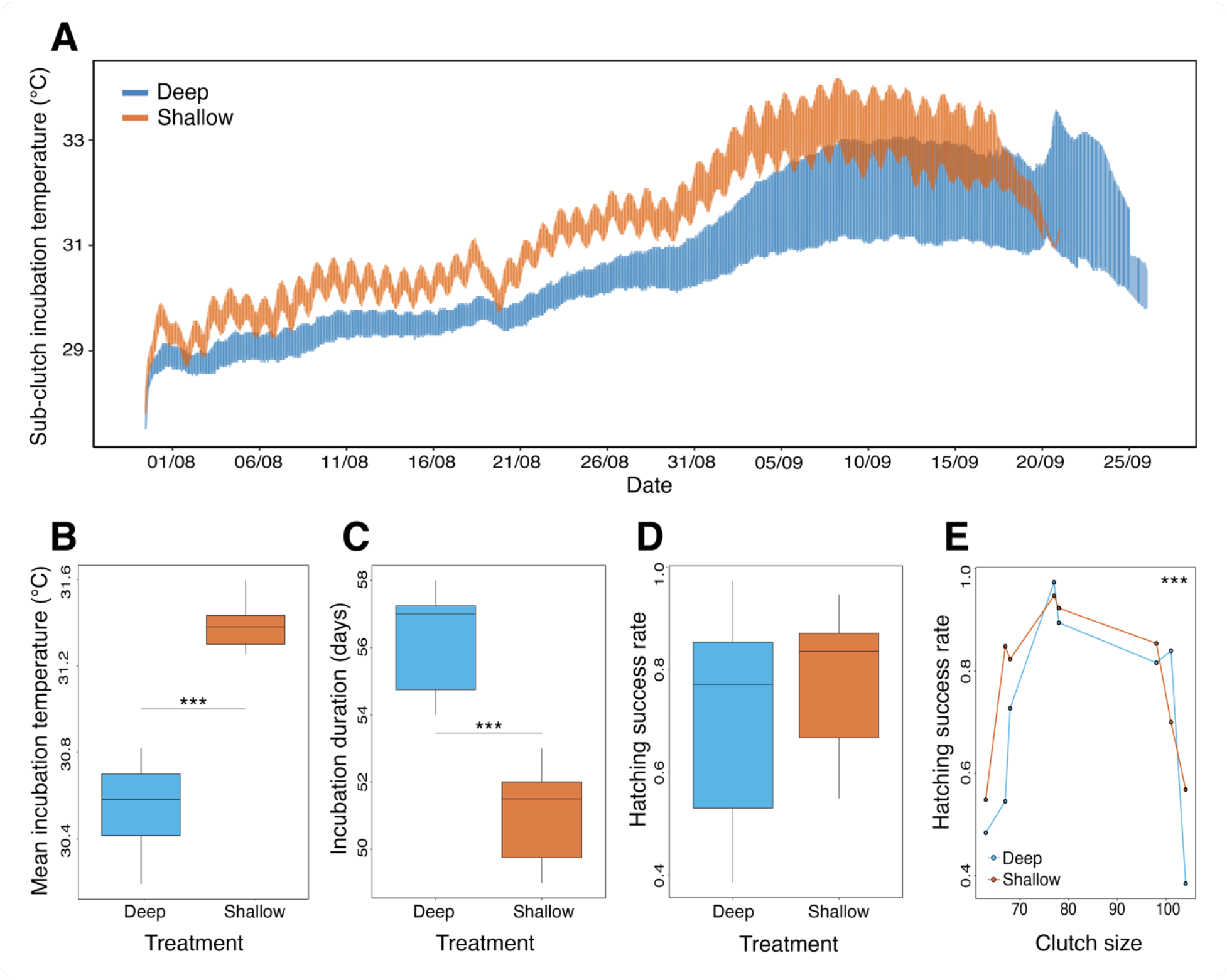
Clutch-level incubation temperature and phenotypes. Deep sub-clutches are coloured in blue and shallow sub-clutches in orange. **(A)** Temperature readings (°C) overlapped across all temperature loggers (n=16) over the entire incubation period. Shallow sub-clutches reached higher temperatures than deep sub-clutches throughout the incubation period, with daily fluctuations visible. **(B)** Sub-clutches incubated in the shallow treatment exhibited higher mean incubation temperatures (°C) than in the deep treatment. **(C)** Incubation duration (days) was longer in sub-clutches incubated in the deep treatment than in the shallow treatment. **(D)** No difference in hatching success rate was found between sub-clutches incubated in deep and shallow treatments. **(E)** Large and small total clutch sizes exhibited lower hatching success rate compared to intermediate clutch sizes.

To confirm the depth treatment was associated with different temperatures, we compared the mean incubation temperature between sub-clutches buried at deep and shallow depths with temperature loggers (16 sub-clutches in total, 8 sub-clutches in each depth treatment). Shallow sub-clutches were confirmed to be warmer than deep sub-clutches (F_1,14_=100.69, p<0.001) by a mean of 0.83°C (**Figure 1B**). Shallow sub-clutches also experienced higher maximum temperatures (T_deep_= 32.64°C ± 0.67°C, T_shallow_= 33.77°C ± 0.40°C, Model statistics: F_1,14_=16.64, p=0.001) and increased temperature fluctuations (mean standard deviation, SD_deep_=1.24°C; SD_shallow_=1.42°C, F=2.91, p=0.11) compared to deep sub-clutches. Hence, the shallow treatment matches predicted conditions of increased temperatures and variability before the end of the century (Laloë et al. 2014).

Increased temperatures translated into faster egg development and shorter incubation duration for the shallow sub-clutches (F_1,14_=43.48, p<0.001, **Figure 1C**). Shallow sub-clutches hatched on average 5 days faster than deep sub-clutches (shallow sub-clutches = 50.7 ± 1.67 days, deep sub-clutches = 56.2 ± 1.51 days). There was no significant difference in hatching success rate of sub-clutches based on depth treatment (F_1,9_=1.764, p=0.217, **Figure 1D**) or mean incubation temperature (F_1,9_=1.885, p= 0.203). Interestingly, we instead found that hatching success was a trait correlated with the total clutch size, with small and large clutch sizes exhibiting lower hatching success rate than intermediate sizes (quadratic term for clutch size, F_1,9_=33.30, p<0.001; Clutch size^2^ x Treatment, F=0.568, p=0.470, **Figure 1E**). Noteworthy, we could detect the effects of metabolic heat, as demonstrated by the positive correlation between incubation temperature and nest size, independently of the depth treatment (F_1,12_=23.036, p<0.001, **Figure S1B**).

### 2. Hatchling-level morphometric and locomotion phenotypes

Across all 10 clutches, we sampled 408 hatchlings (n=198 from 20 deep sub-clutches, n=210 from 20 shallow sub-clutches) for morphometrics and fitness tests. As expected, there was a positive correlation between hatchling SCL and mass (F=299.091, p<0.001), and this correlation varied with depth treatment (Mass x Treatment, F=9.746, p=0.002, **Figure 2A**). We found that hatchling SCL also correlated with the interaction between clutch size and mean sub-clutch incubation temperature (F=9.832, p=0.002), whereby smaller clutch sizes are associated with increased SCL, but this link is altered by sub-clutch temperature.

**Figure 2.**
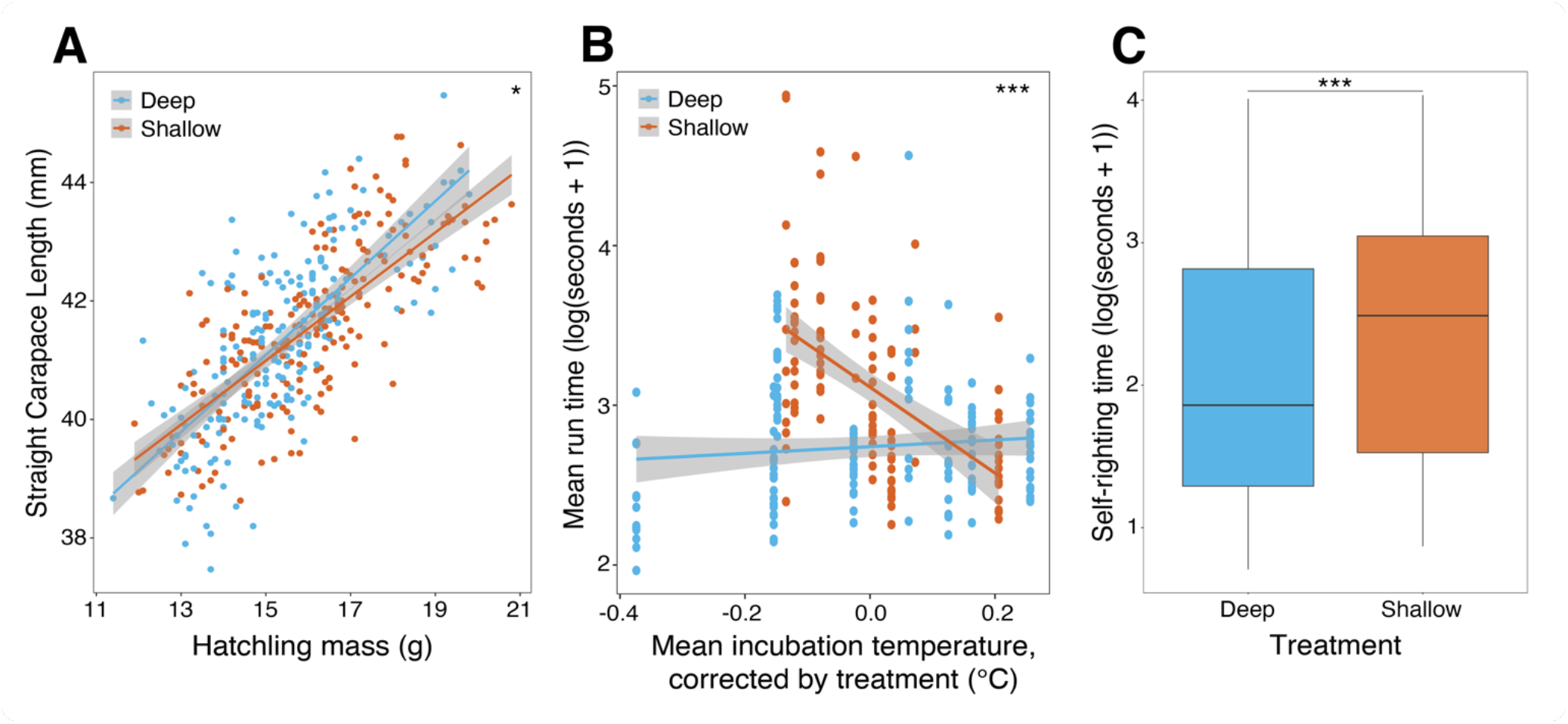
Individual-level, hatchling morphometric and locomotion phenotypes. Hatchlings emerging from deep sub-clutches are shown in blue and those from shallow sub-clutches in orange. **(A)** Hatchling straight carapace length (SCL, mm) is positively correlated with their mass (g), but this allometric link is altered by the depth incubation treatment. **(B)** Relationship between the time taken to run 50 cm (log(seconds+1)) and the interaction between incubation temperature (°C) and the depth treatment, shown in residuals. **(C)** Time taken to self-right (log(seconds+1)) is slower for hatchlings that emerged from sub-clutches incubated in the shallow treatment.

To go beyond morphological traits, we also measured the locomotion capacity of hatchlings, which is essential for dispersal and predator avoidance immediately upon emergence. Mean run time was correlated with an interaction between depth treatment and mean incubation temperature (F=20.093, p<0.0001, **Figure 2B**). We further found that for a given clutch size, hatchlings from the deep sub-clutches consistently outperformed hatchling from shallow sub-clutches, particularly in lower clutch sizes (F=4.568, p =0.033). There was no correlation between self-righting time and mean incubation temperature, clutch size or hatchling size, which were all removed from the final model. Instead, the best predictor of self-righting was the depth treatment, with hatchling emerging from the shallow sub-clutches being slower at self-righting (F= 11.489, p<0.001, **Figure 2C**).

### 3. Global DNA methylation patterns

For a subset of 40 hatchlings (n=20 per depth treatment, n=2 per sub-clutch), we collected red blood cell samples immediately after fitness tests for WGBS. This enabled us to investigate how global DNA methylation patterns are impacted by incubation condition on a genome-wide scale, using our dataset of 2,733,573 CpG sites that were covered in all individuals. These CpG sites were distributed across gene promoters (n=45,627, 1.67%), exons (n=73,374, 2.68%), introns (n=1,245,062, 45.55%) and intergenic space (n=1,369,510, 50.1%) throughout the genome (**Figure 3A**).

**Figure 3.**
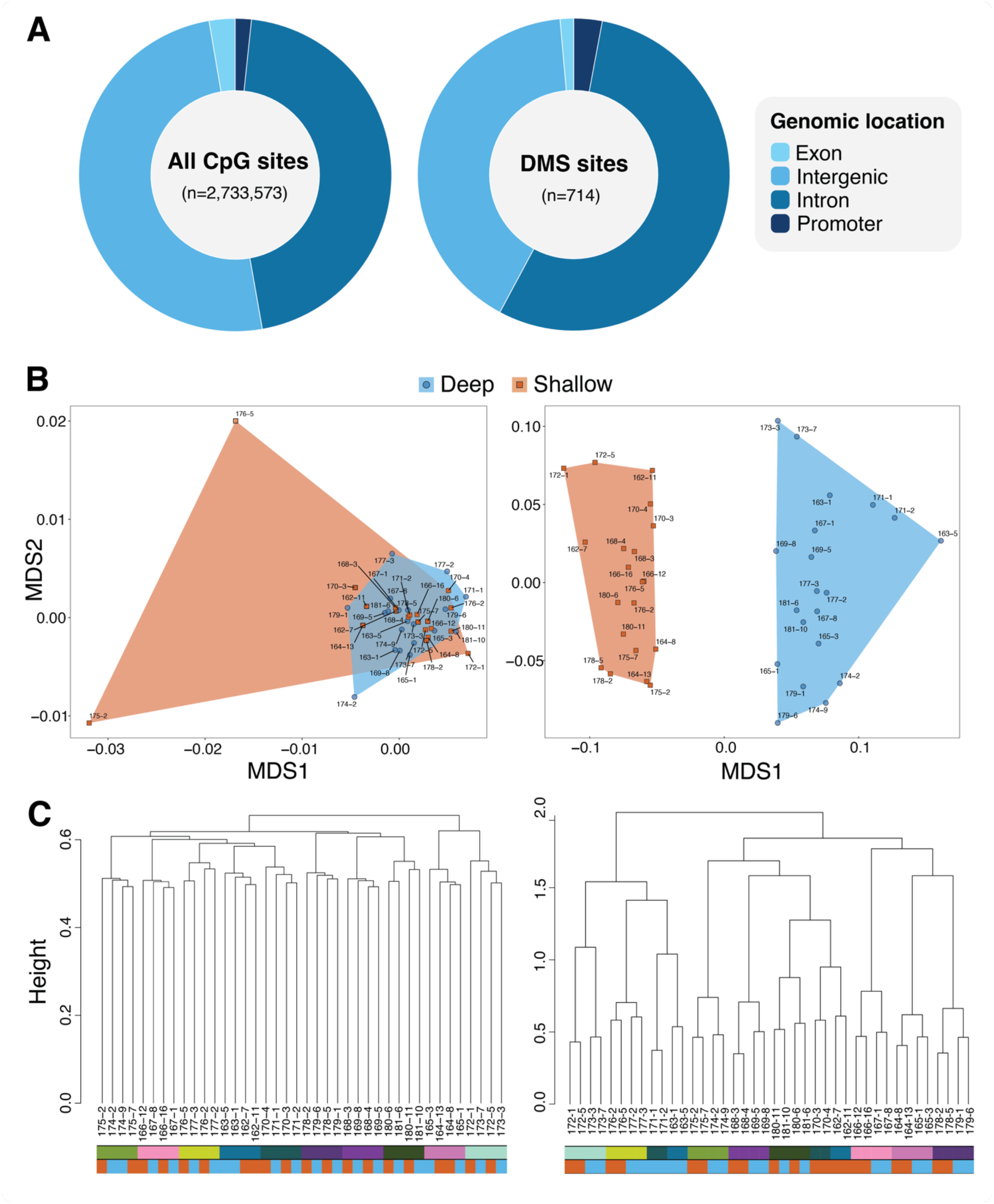
DNA methylation patterns of 40 hatchlings sequenced via WGBS. All plots on the left-hand side show global methylation (n=2,733,573 CpG sites) results, and all plots on the right-hand side show results for DMS (n=714 CpG sites). **(A)** Proportions of CpG sites across four genomic feature types. Left: global CpG sites, on gene promoters (1.67%), exons (2.68%), introns (45.55%) and intergenic space (50.1%). Right: DMS, on gene promoters (2.94%), exons (1.4%), introns (54.9%) and intergenic regions (40.76%). CpG sites were similarly distributed across genomic features between global and DMS categories. **(B)** NMDS plots with hatchlings coloured by treatment (shallow in orange, deep in blue). Left: global methylation. Right: DMS. Clear clustering by incubation treatment can be seen in the DMS but not in global methylation plot. **(C)** Hierarchical clustering dendrograms, with colour bars underneath the dendrogram representing each hatchling’s maternal ID (top bar) and depth treatment (bottom bar, with shallow in orange, deep in blue). Left: global methylation. Right: DMS. Maternal ID best explains the clustering in both global methylation and DMS, with some clustering signal by depth treatment for DMS.

NMDS and PERMANOVA analyses showed that hatchlings did not form separate clusters by depth treatment, based on their global DNA methylation values (**Figure 3B, Figure S3A**, R^2^=0.024, F=0.98, p=0.80). These results were robust even after the exclusion of two potential outliers that passed all quality controls (sample IDs: 175-2 and 176-5, **Figure S3A** for NMDS plots produced without these samples). Instead, we showed that maternal ID best explained the clustering of global methylation variance (R^2^=0.25, F=1.11, p=0.001). This result was confirmed by a hierarchical clustering analysis, and hence highlights the important of family genetic background in driving DNA methylation patterns (**Figure 3C**). The lack of an important role for depth treatment in determining global methylation is further supported by our finding that total count of methylated CpG sites per individual does not differ between hatchling incubated at different depth treatments, in either the count of sites with non-zero methylation (F=0.041, p=0.84, **Figure S2A**) or sites with >70% methylation (F=0.115, p=0.74, **Figure S2B**).

### 4. DMS identified between incubation treatments

Whilst incubation treatment did not leave a detectable molecular signature at the whole methylome-level, we identified 714 DMS between hatchlings from different depth treatments, once corrected for maternal ID as a covariate (methylation difference >15%, q-value <0.01, **Figure S4**). Mean methylation differences ranged from 38.1% hyper-methylated (n=318) in hatchling from the cooler, deep treatment, to 34.0% hyper-methylated (n=396) in hatchlings from the warmer, shallow treatment (i.e. equivalent to hypo-methylated against hatchlings from the deep treatment, **Figure S5**). These DMS were located across gene promoters (n=21, 2.94%), exons (n=10, 1.4%), introns (n=392, 54.9%) and intergenic regions (n=291, 40.76%), which were similarly distributed across those genomic features compared to global CpG sites (χ^2^=2.58, p=0.46, **Figure 3A**).

Hatchling methylation values at these 714 DMS generated clear clustering by depth treatments in the NMDS plot (**Figure 3B**). A PERMANOVA showed that clustering at DMS is associated to the depth treatment (R^2^=0.15, F=10.55, p=0.001) in addition to maternal ID (R^2^=0.44, F=3.48, p=0.001), but not to their interaction. The effect of maternal genetic origin can also be observed in the hierarchical clustering analyses, despite some structuring by the depth treatment (**Figure 3C**). This again highlights the importance of genetic background on DNA methylation patterning. Overall, methylation variance at these 714 DMS strongly discriminated between hatchlings that experienced the cooler, deep treatment and warmer, shallow treatment.

### 5. Genes associated with DMS

From the 714 DMS, 483 DMS were associated to a gene via overlap or proximity to the TSS (<10 kb away) if it was intergenic **(Table S2).** Of these, 223 sites were hyper-methylated in deep-incubated hatchlings, versus 260 sites being hyper-methylated in shallow-incubated hatchlings **(Figure S5).** Two genes had four DMS, two genes had three DMS, 33 genes had two DMS and 403 genes had one DMS located on them. Given that differentially methylated CpG sites tend to co-occur in clusters in the genome (Suzuki and Bird, 2008), the small number of genes with multiple DMS in our dataset is likely due to our stringent filtering for sites covered in all individuals, as needed to start identifying biomarkers.

We further investigated the functions of genes associated with the 21 DMS of interest that met our selection criteria (**Figure 4A, Table S3**). On chromosome 11, we identified RBMS1 (RNA Binding Motif Single Stranded Interacting Protein 1) as one of the genes with the highest number of DMS, with four DMS all found on an intron (DMS ID: RMBS1.1, RMBS1.2, RMBS1.3, RMBS1.4). This gene encodes an RNA-binding protein involved in endocytosis, vesicle trafficking and cytoskeletal regulation. Interestingly, two DMS were more methylated in deep-incubated hatchlings (mean difference = 30.6%), whereas the other two DMS were more methylated in shallow-incubated hatchlings (mean difference = −26.4%). The other gene with four DMS (DMS ID: TEX14.1, TEX14.2, TEX14.3, TEX14.4; mean difference=20.0%) was identified as TEX14 (Testis Expressed 14, Intercellular Bridge Forming Factor) on chromosome 17. This gene encodes a protein necessary for intercellular bridges in germ cells, which are required for spermatogenesis. All DMS on this gene exhibited higher methylation values in deep-incubated hatchlings and were also located on an intron.

**Figure 4.**
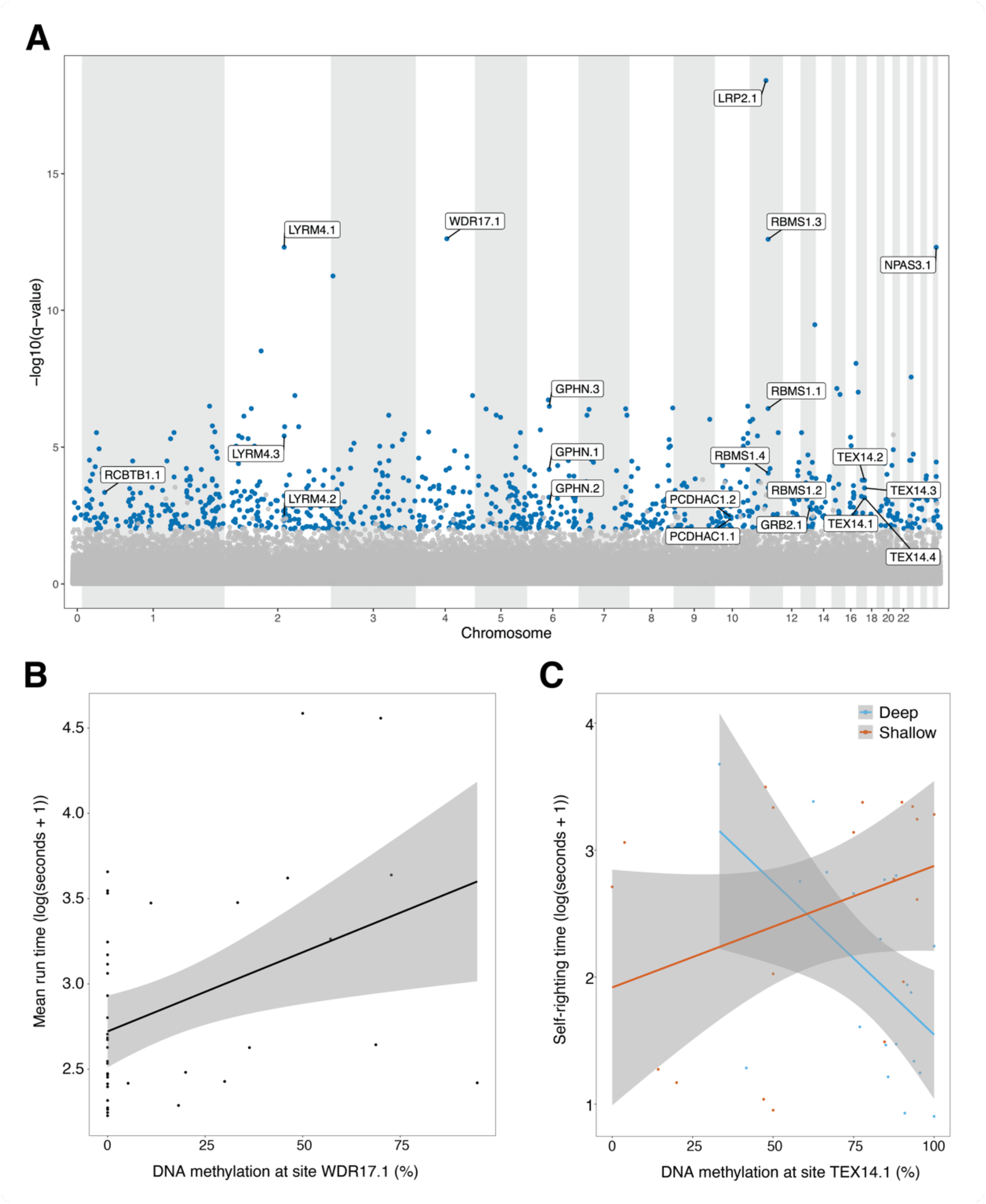
Methylation results in relation to DMS of interest. **(A)** Manhattan plot of the location of all CpG sites (n=2,733,573) across the genome against the –log_10_(q-value), with all DMS (n=714, methylation difference >15%, q-value < 0.01) coloured in blue, and the 21 DMS of interest labelled. **(B)** Positive relationship between DNA methylation value (%) per individual at the DMS site on the WDR17 gene (WDR17.1) against the mean run time (log(seconds+1)). **(C)** Relationship between DNA methylation value (%) per individual at the first DMS site on the TEX14 gene (TEX14.1) against the mean self-righting time (log(seconds+1)). Self-righting time exhibited an interaction between depth treatment and methylation value at this site, with faster self-righting times and higher methylation values in hatchlings from deep sub-clutches, and the opposite case for hatchlings from shallow sub-clutches.

Two other genes had three DMS each, all on introns. The three DMS located on LYRM4 (LYR Motif Containing 4) showed higher methylation in shallow-incubated hatchlings (DMS ID: LYRM4.1, LYRM4.2, LYRM4.3; mean difference=-25.9%). The GPHN (Gephyrin) gene however had one DMS that was more methylated in shallow-incubated hatchlings (DMS ID: GPHN1.1; difference=-23.5%) and two DMS that were more methylated in deep-incubated hatchlings (DMS ID: GPHN1.2, GPHN1.3; mean difference=16.8%). Whilst LYRM4 encodes a protein linked to cytosolic iron homeostasis that is found in both the mitochondria and nucleus, GPHN encodes for a neuronal assembly protein that anchors inhibitory neurotransmitter receptors to the postsynaptic cytoskeleton.

After identifying genes with the most DMS, we next looked at genes associated with the top five most differentiated DMS on the Manhattan plot (**Figure 4A**). This criterion already identified DMS on RBMS1 (DMS ID: RBMS1.3, methylation difference=38.1%) and LYRM4 (DMS ID: LYRM4.1, methylation difference=-34.1%). The other three DMS were all located on introns of different genes. On LRP2 (LDL receptor related protein 2), the DMS was more methylated in deep-incubated hatchlings (DMS ID: LRP2.1, difference= 24.4%). This gene encodes an endocytic receptor primarily expressed in epithelial tissues, with various functions including immune responses, vitamin absorption, lipid transport in the bloodstream, stress responses and development. The remaining two most differentiated DMS are located on the WDR17 (WD Repeat Domain 17) and NPAS3 (Neuronal PAS Domain Protein) genes, which were both more methylated in shallow-incubated hatchlings (DMS ID: WDR17.1; difference=-25.8%, DMS ID: NPAS3.1; difference=-31.5%). The protein encoded of NPAS3 is localized to the nucleus and regulates genes involved in neurogenesis, meanwhile WDR17 is expressed in the retina and testis, but the function of its protein remains to be understood.

Finally, we focused on DMS associated with promoters, as DNA methylation regulation at promoter regions is best characterised to date (Jones, 2012). The gene with the most DMS associated to it was PCDHAC1, with two DMS that have higher methylation in deep-incubated hatchlings (PCDHAC1.1, PCDHAC1.2; mean difference=19.0%). This gene is involved in synaptic adhesion and specific cell-cell connections in the brain. The most differentiated DMS on promoters were located on the GRB2 (Growth Factor Receptor Bound Protein 2) and RCBTB1 (RCC1 And BTB Domain Containing Protein 1) genes, which were more methylated in deep-incubated hatchlings (GRB2.1; difference= 22.9%) and shallow-incubated hatchlings (RCBTB1.1; difference=-24.5%) respectively. GRB2 encodes an epidermal adaptor protein that participates in signal transduction pathways, whilst RCBTB1 induces cellular hypertrophy when over-expressed in vascular smooth muscle cells of rats.

### 6. Correlating methylation value at DMS of interest against fitness-related phenotypes

We further tested the relationship between individual hatchling fitness-related traits and methylation value at the 21 DMS of interest. We created linear mixed effects models for each of the fitness traits (SCL, mass, average run length, average self-righting time), with all models containing maternal ID as a random effect. Methylation values, depth treatment and their interaction were used as fixed factors. After correcting for multiple testing, we found that mean run time over 50cm was correlated with methylation values of DMS at the WDR17 gene (t=2.876, p=0.0067; **Figure 4B**), with higher methylation associated with longer crawl times. We also found self-righting time was explained by an interaction between depth treatment and DNA methylation at a site on the TEX14 gene (t=2.923, p=0.006; **Figure 4C**). Higher methylation values were associated with lower self-righting time in hatchling from deep nests, whereas the opposite was true for hatchling from shallow nests.

### 7. Functional enrichment of GO terms

To investigate the functions of identified DMS more broadly than our manually selected 21 DMS of interest, we performed a GO term enrichment analysis. We identified 87 enrichments for biological processes, of which 48 were linked to hypermethylation in deep-incubated hatchlings and 39 in shallow-incubated hatchlings (**Figure 5, Table S4)**. The only enriched process shared between both methylation directions was the molybdopterin cofactor biosynthetic process. This cofactor is involved in DNA regulation via DNA methylation and gene expression (Mendel 2013). It is also known for its role in development and growth, where molybdoenzymes help metabolize RNA and are important for organ development (Schwarz et al. 2009). Enrichment of cellular component terms were more prevalent in hatchlings from the shallow treatment with 25 terms, versus five terms for hatchlings in their deep counterparts. The top GO term in shallow-incubated hatchlings related to the mitochondrial intermembrane space, which provides a regulatory compartment that influences respiratory function, cell death pathways, and redox signalling (Kühlbrandt 2015). Dysregulation of respiratory functions taking place in the intermembrane space can alter the processes of oxidative phosphorylation and ATP production. Given that hatchlings undergo a highly active behavioural stage upon emergence, known as the swimming frenzy, disruption to such energy-related processes may hinder active dispersal (e.g. Jones et al 2007). In relation to molecular functions, 18 terms were enriched in shallow-incubated hatchlings, against 16 terms in deep-incubated hatchlings. Most functions hyper-methylated in shallow-incubated hatchlings were linked to cellular homeostasis, such as pH and redox state regulation, and cell proliferation signalling pathways. For functions hyper-methylated in shallow-incubated hatchlings, the enriched molecular functions suggested a link to cytoskeleton regulation. Indeed, myosin binding proteins interact with myosin motors composed of heavy and light chains that move along actin filaments. 1-phosphatidylinositol helps anchor the cytoskeleton to membranes. So, these interactions likely cooperate to regulate cytoskeletal dynamics (Saarikangas et al 2010).

**Figure 5.**
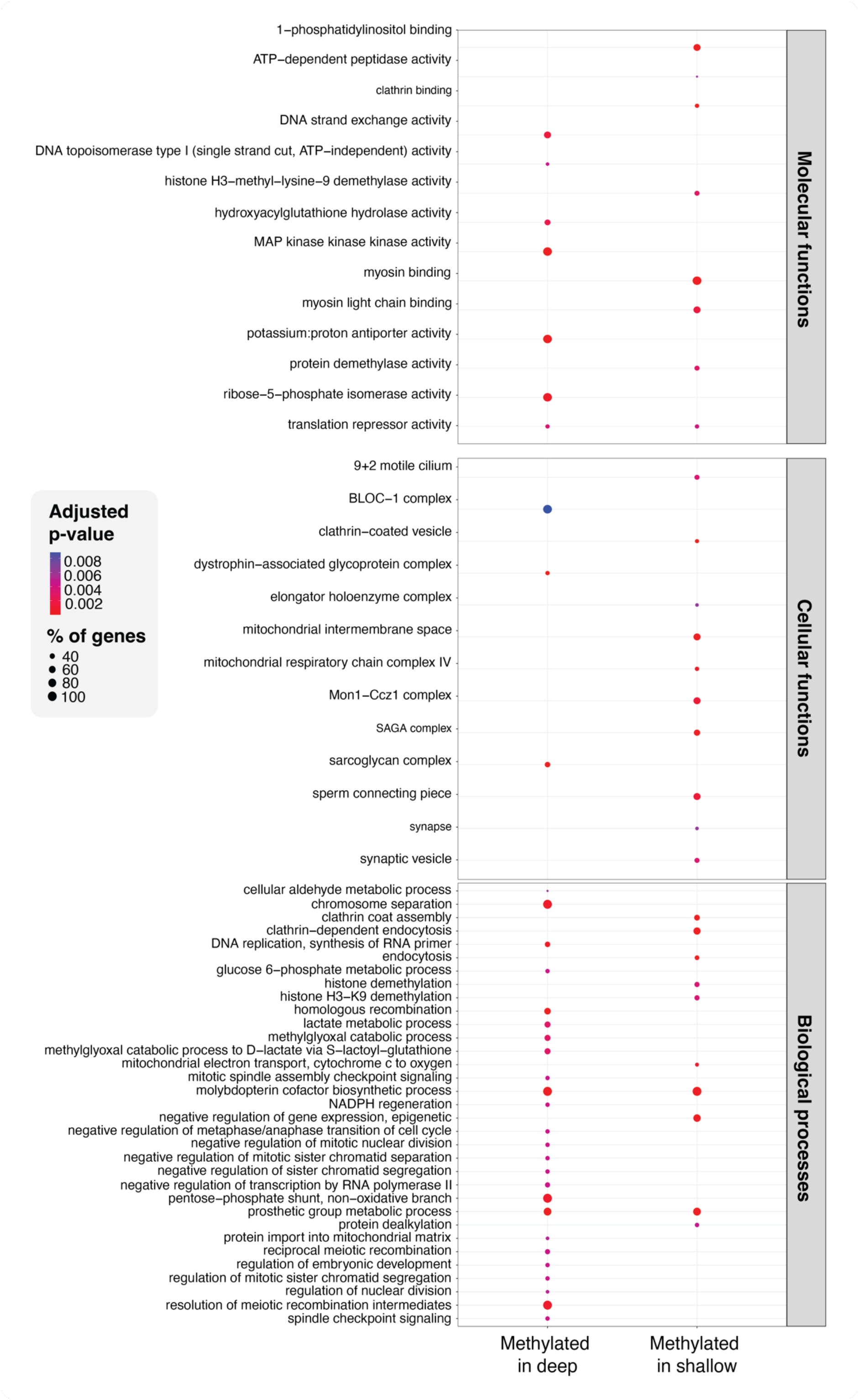
Functional enrichment of GO terms associated with genes that exhibit DMS in hatchlings from the incubation treatments. Sites with higher methylation values in hatchlings from the deep treatment are shown on the left, and while sites with higher methylation values in hatchlings from the shallow treatment are shown on the right. The top panel shows GO terms related to molecular functions, the middle panel shows GO terms related to cellular functions, and the bottom panel shows GO terms related to biological processes. Dot size represents the percentage of genes of a given term enriched in our dataset. The colour scale relates to the adjusted p-values. Only GO terms that were present in at least 25% of genes are shown in this figure.

## Discussion

As climate change intensifies, the proximal, thermal responses of individuals will be physiological and likely sublethal. Monitoring these sublethal effects in threatened species is essential, but reliable and high-throughput molecular markers are needed. By experimentally manipulating incubation temperatures to simulate predicted future conditions, we found that warmer temperatures altered the length-mass relationship of hatchlings and reduced their crawling speed and self-righting ability, despite high and comparable hatching success rates between incubation treatments. Next, using whole-genome bisulfite sequencing, we found 714 CpG sites that were consistently differentially methylated between hatchlings from the deep, cooler and shallow, warmer treatments. This demonstrates that incubation temperature leaves detectable physiological signatures in the blood methylomes of loggerhead sea turtle hatchlings. Interestingly, we found that multiple DMS occurred on genes with functions in neurodevelopment. Overall, our results show that even when sea turtles survive high incubation temperatures, they can still suffer negative fitness consequences in key physiological traits that impact dispersal ability. Our study also brings evidence that DNA methylation can be used as potential biomarkers of early life thermal stress and their associated sublethal effects.

Our findings reveal significant impacts of thermal incubation regime on important fitness-related phenotypes of hatchlings, in particular their length-mass relationship and their crawling and self-righting capacity. The two measured locomotion traits were slower in hatchlings incubated at the elevated temperatures of the shallow treatment. Our results align with previous suggestions of reduced fitness in loggerhead sea turtles under higher incubation temperatures (e.g. Fisher et al 2014; Booth, 2017; Reece et al. 2002, Fleming et al. 2020). Crawling and self-righting are part of the terrestrial dispersal phase immediately after emergence, which is followed by the swimming frenzy phase in the ocean (Scott et al. 2014). If temperature persistently hampers hatchling locomotion, it could impede their ability to disperse and colonize new habitats as climate change shifts nesting distributions (Perry et al 2005; Kobayashi et al 2018; Duffy et al. 2022). Reduced locomotion will also weaken their predator avoidance ability and secondarily their survival, which could negatively impact population recruitment rates (Hamilton et al. 2016). Furthermore, smaller hatchlings already exhibit reduced swimming capacity compared to larger counterparts (Mueller et al. 2019; Scott et al. 2014), so compounding effects on dispersal may be underestimated. If these impacts continue beyond the dispersal stage, carryover effects across life stages could also be underestimated. Noteworthy, numerous studies have shown similar effects of incubation temperature on other physiological and performance traits in other sea turtle species (e.g. Howard et al. 2014; Gatto et al., 2022).

To evaluate the effects of temperature and incubation conditions on hatchlings, we used a split clutch-design in an *in-situ* hatchery. This experimental approach is valuable for assessing environmental impacts on species, independently of their genetic background. Whilst extensively used in laboratory settings (e.g. Eizaguirre et al. 2012, Kaufman et al. 2014), such approaches are often constrained in field conditions. In the context of sea turtles, hatchery experiments have often focused on management and reduction of incubation temperatures to mitigate lethal effects of increasing temperatures (e.g. Clarke et al 2021; Esteban et al. 2018; Yao et al. 2022). Here, the experimental nests did not approach the lethal limits of eggs and embryos, as indicated by high, comparable hatching success rates between sub-clutches incubated across both depth treatments. Instead, it helped revealed that hatching success rate may be a clutch-specific trait independent of incubation conditions at sublethal thermal ranges, where small and large clutch sizes had increased failures compared to intermediate clutch sizes.

By pairing our findings of phenotypic fitness reductions with the functional annotation of differentially methylated genes, we bring physiological and molecular evidence for important sublethal effects. This study adds 714 differentially methylated sites to growing evidence that DNA methylation carries the signatures of temperature exposure, even within sublethal ranges (Le Luyer et al. 2017; Sheldon et al. 2020; Bock et al. 2020). Gene functions associated with identified DMS suggest wide-ranging impacts on neurodevelopmental processes. For instance, the DMS on the NPAS3 gene, which showed reduced methylation in deep-incubated hatchlings, regulates neurogenesis and has been associated with neurodevelopmental disorders when dysregulated (Kamm et al. 2013; Yang et al. 2016). Similarly, DMS were detected on GPHN (gephyrin). This gene is essential for synaptic function (Ramming et al. 2000), and mutations in humans are associated with neurodevelopmental disorders (Lionel et al. 2013). Beyond specific genes, the breadth of affected pathways and functions included cytoskeletal dynamics and cellular adhesion, highlighting the potential for pervasive developmental disruption from even small temperature shifts, as seen in other vertebrates (Metzger and Schulte 2017; Anastasiadi et al. 2021). Sublethal disruption of these diverse functions could explain the observed fitness reductions in shallow-incubated hatchlings, with some direct links indeed found between locomotion capacity and DNA methylation status on two genes.

As expected from incubating the eggs of a TSD species under different temperatures, we detected differential methylation at genes involved in gonadal development. Specifically, TEX14 encodes a protein that localizes to germ cell intercellular bridges, essential for spermatogenesis and fertility (Greenbaum et al. 2009). DNA methylation of TEX14 was associated with self-righting time in this study in a treatment-dependent manner. This offers a candidate gene to test the predictions of the Charnov-Bull model of TSD evolution at the molecular level (Charnov and Bull, 1977). This evolutionary model predicts sex-specific fitness advantages are associated with sex-specific thermal environments, which should have a molecular underpinning.

Taken together, the 714 DMS identified in this study could form a set of potential epigenetic biomarkers to monitor the sublethal effects of incubation temperatures. This is strengthened by the strict thresholds we applied, whereby CpG sites had to be identified as differentially methylated across all hatchlings. Similar signatures of incubation stress have been found in coho salmon (*Oncorhynchus kisutch*). In hatchery-reared versus wild individuals of this species, a large proportion of methylation variation was associated with rearing environment, with enrichment for biological functions broadly correlated with the capacity of hatchery-born smolts to migrate successfully to the ocean (Le Luyer et al. 2017). It was further shown that those early-life methylation marks persisted into germ line cells (Leitwein et al 2021). If this is also true for sea turtles, it opens the possibility for detecting signature of incubation condition at later life stages of this long-lived species.

Of note, the main clustering determinant at both global and differentially methylated CpG sites was the maternal ID, which represents the genetic background of the clutches. This supports a growing body of research documenting the strong influence of genetic background on DNA methylation (Yang et al. 2010; Rey et al. 2020). For example, Sepers et al. (2023) showed that genetic background was a better predictor of DNA methylation than rearing environment in a wild great tit (*Parus major*) population. Genetic sequence also explained variation in DNA methylation across 580 animal species, with the density of CpG sites and islands thought to be particularly important (Klughammer et al. 2023). As molecular markers for monitoring responses to climate change are being developed, combining genetic and epigenetic data to disentangle the effects of environment versus genetics on methylation patterns will provide crucial insights.

Beyond sea turtles, the framework we introduce here - leveraging split-clutch experimental designs to identify epigenetic biomarkers associated with climate-mediated stressors in the field - could be widely applicable to conservation biology questions. Whilst many laboratory and captive studies have documented thermal effects on epigenetic patterns, demonstrating these relationships in natural contexts remains challenging. Manipulative field studies are thus invaluable, yet still rare in ecological epigenetics, especially in non-model species (Chapelle and Silvestre, 2022). Our findings showcase their utility for strengthening claims of association between environmental stressors and epigenetic variation. Expanding the epigenetic toolkit to wild populations could similarly reveal subtle impacts of climate threats that evade detection by traditional survival-oriented metrics. Well-designed field studies in non-model systems will further provide the most realistic insights into the interplay between phenotypic fitness, epigenetic regulation and population resilience across ecological contexts in the wild, which is a priority for advancing predictive, mechanistic understanding of conservation epigenetics.

## Conclusions

Our findings reveal that beyond mortality, warming could have pervasive yet cryptic sublethal impacts in sea turtles. Even if lethal thresholds are not crossed, subtler fitness reductions could still accelerate population declines, especially if impacts in early life are carried over to later life stages. Such impacts are however detectable through DNA methylation status at identified genomic sites. As these molecular markers carry signatures of thermal stress induced by the incubation environment, they could provide a means to monitor early warning signals of sublethal effects at high temperatures, complementing demographic perspectives. Our study also highlights the complex interplay between genetics and environment that will shape population trajectories and therefore species persistence in a changing world. Whilst much work remains to unravel these dynamics in non-model systems, emerging conservation epigenetic approaches offer promising tools to meet the challenges ahead.

## Data Availability

All data and scripts will be made available after the manuscript is accepted for publication.

## Permits

All sample collection and experiments adhered to national legislation and were approved by the Direção Nacional do Ambiente of Cabo Verde (authorisation: 037/DNA/2021).

## Author Contributions

C.E and E.C.Y designed the study with support of S.M.C., S.J.R. and J.M.M-D. E.C.Y, I.O.A, C.E, K.F, D.N and A.T performed fieldwork, with coordination by A.L, K.F, A.T, S.M.C and C.E. E.C.Y performed DNA extractions. C.E and J.D.G performed clutch and hatchling fitness analyses. E.C.Y performed read quality control and methylation calling. J.D.G performed SNP calling. E.C.Y performed methylation analyses, functional annotation and enrichment analyses, with input and scripts adapted from A.B. E.C.Y, C.E and J.D.G wrote the manuscript. All authors contributed to the final version of the manuscript.

## Acknowledgements

The authors would like to thank Carla Moniz, Emma Agnelli and volunteers with Project Biodiversity for their support in the hatchery.

## List of Supplementary Material

**Supplementary Figure 1.** Relationships of mean incubation temperature with incubation duration and clutch size

**Supplementary Figure 2.** Counts of methylated CpG sites per individual

**Supplementary Figure 3.** NMDS plots showing links between 1 to 3 MDS dimensions

**Supplementary Figure 4.** Volcano plot of the 714 DMS identified between incubation treatments

**Supplementary Figure 5.** Manhattan plot of DMS against methylation difference between hatchlings from different incubation treatments

**Supplementary Text 1.** Extended methods for SNP calling pipeline

**Supplementary Table 1.** WGBS statistics per hatchling

**Supplementary Table 2**. Functional annotation of all 483 gene-associated DMS (Excel)

**Supplementary Table 3.** Names and functional annotation of the 21 DMS of interest (Excel)

**Supplementary Table 4.** GO enrichment study results (Excel)

