## Supplementary Material for "DNA methylation carries signatures of sublethal effects under thermal stress in loggerhead sea turtles"

**Supplementary Figure 1.** Relationships of mean incubation temperature with incubation duration and clutch size. Deep sub-clutches are coloured in blue and shallow sub-clutches in orange. **(A)** Correlation between incubation duration (days) and mean sub-clutch incubation temperature (ºC). The total incubation duration is shorter in sub-clutches incubated at higher temperatures. **(B)** Metabolic heat as shown by the positive correlation between incubation temperature and nest size. Note, the interaction between the depth treatment and the clutch size was uncorrelated with metabolic heat, as expected in a split-clutch experimental design (Interaction Treatment by nest size, $F_{1,12}$= 1.3166, p=0.273).

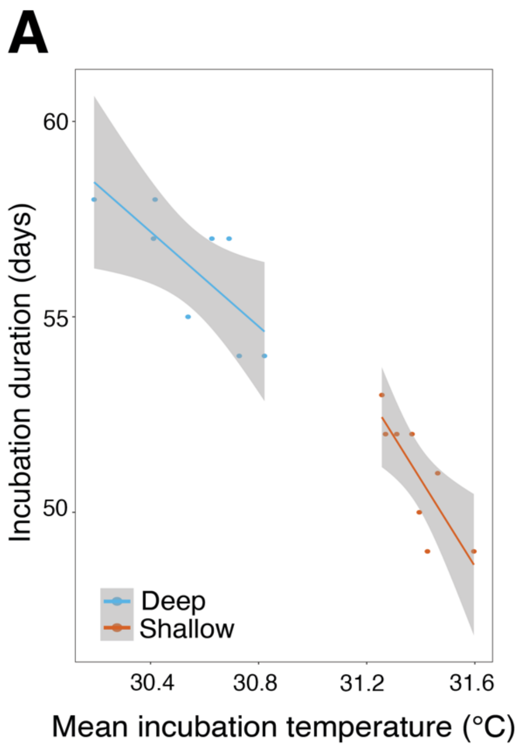

**Supplementary Figure 2.** Counts of methylated CpG sites per individual. Deep-incubated hatchlings are in blue and shallow-incubated hatchlings are in orange. **(A)** Count of CpG sites with non-zero (>0%) methylation values per individual. **(B)** Count of CpG sites with a methylation proportion above 0.7 (>70%) per individual.

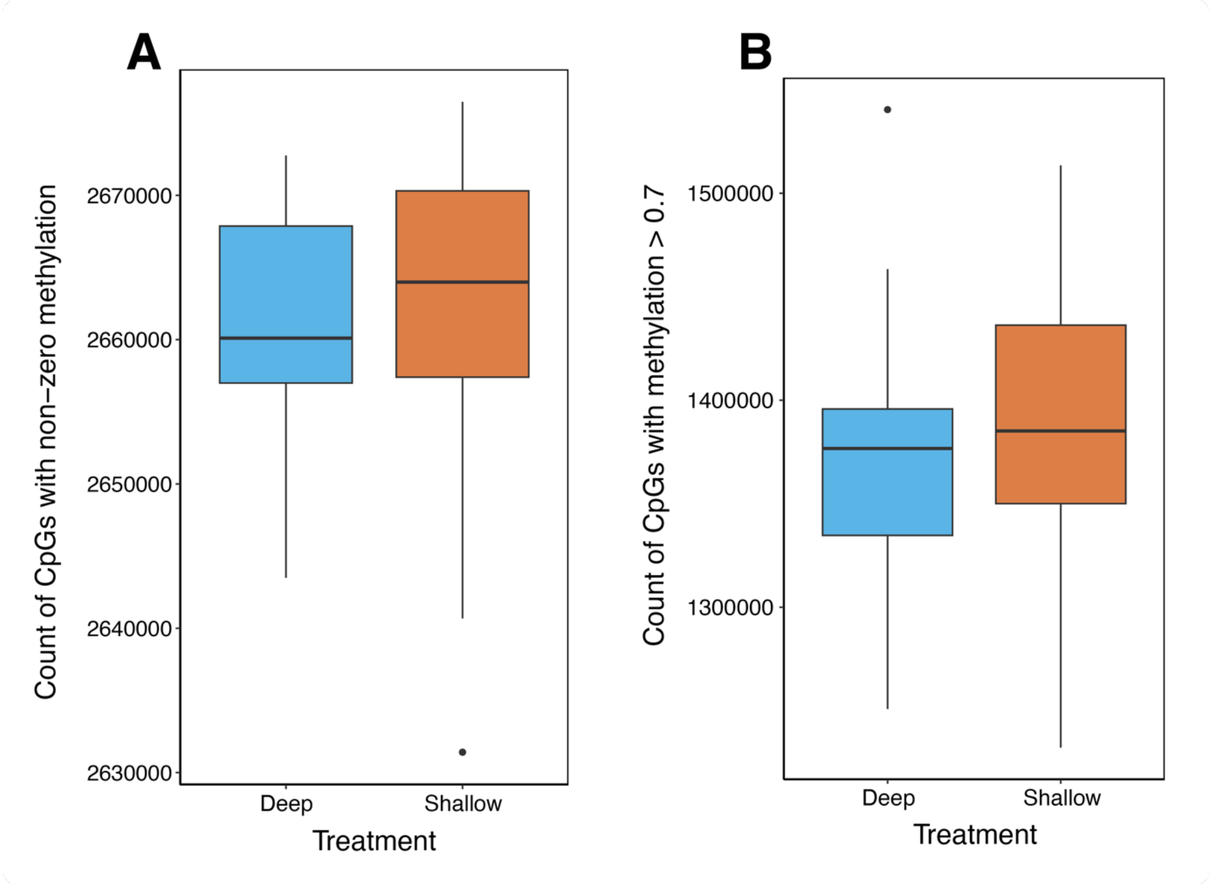

**Supplementary Figure 3.** NMDS plots with additional dimensions. All plots in the left column show global methylation (n=2,733,573 CpG sites) results. All plots in the middle column show global methylation results (n=2,733,573 CpG sites), with two potential outlier individuals excluded (sample IDs: 175-2 and 176-5). All plots in the right column show DMS results (n=714 CpG sites). **(A)** Plots of dimensions MDS1 versus MDS2. **(B)** Plots of dimensions MDS1 versus MDS3. **(C)** Plots of dimensions MDS2 versus MDS3. **(D)** Stress plots of goodness of fit for NMDS analyses.

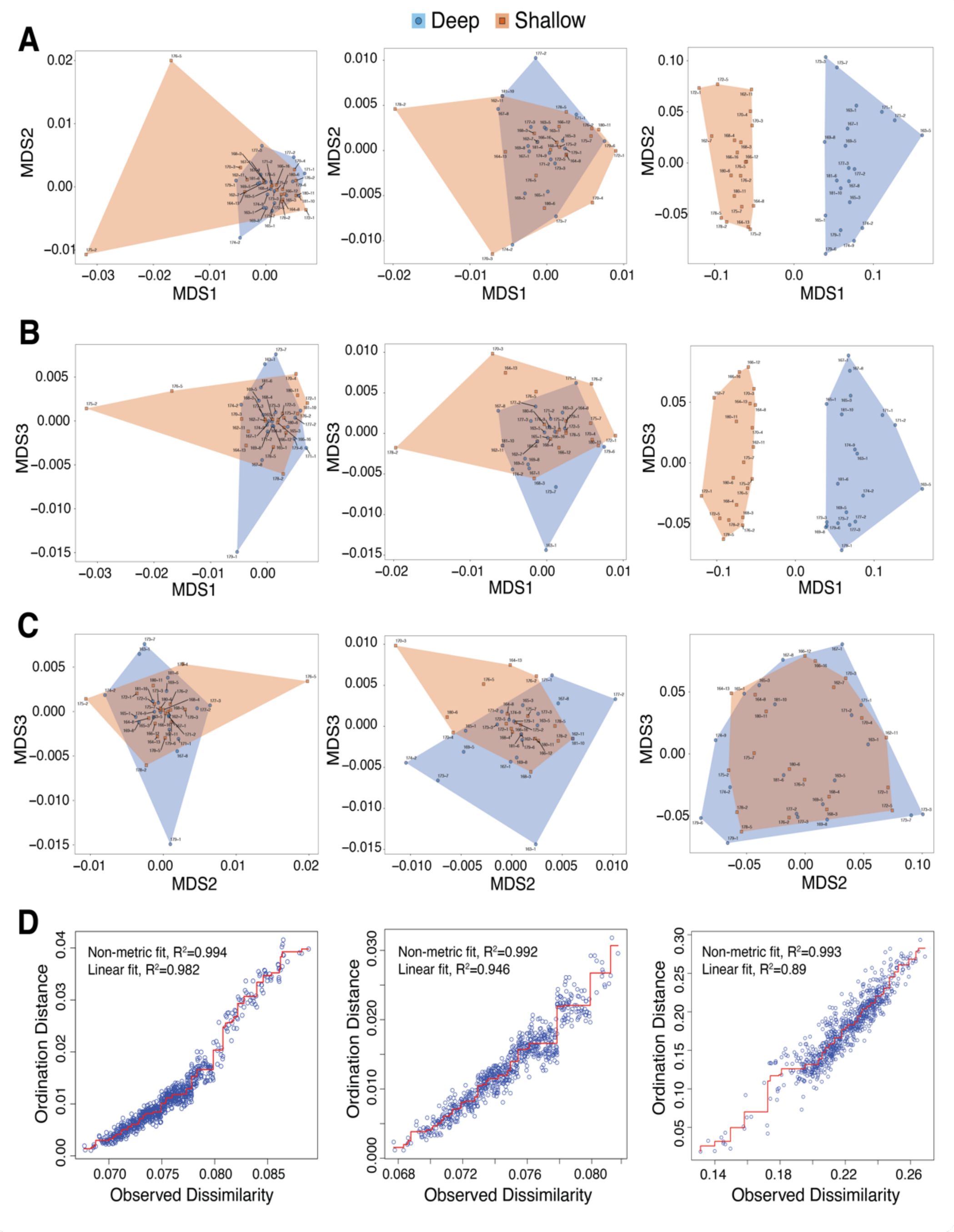

**Supplementary Figure 4.** Volcano plot of the 714 DMS identified between hatchling from the different incubation treatments. DMS were selected with the threshold of having over 15% methylation difference between hatchling from the deep and shallow treatments, and a q-value less than 0.01. For each DMS, its methylation difference is plotted against the $-\log_{10}$(q-value). A positive methylation value indicates hyper-methylation in deep-incubated hatchlings and a negative value indicates hyper-methylation in shallow-incubated hatchlings.

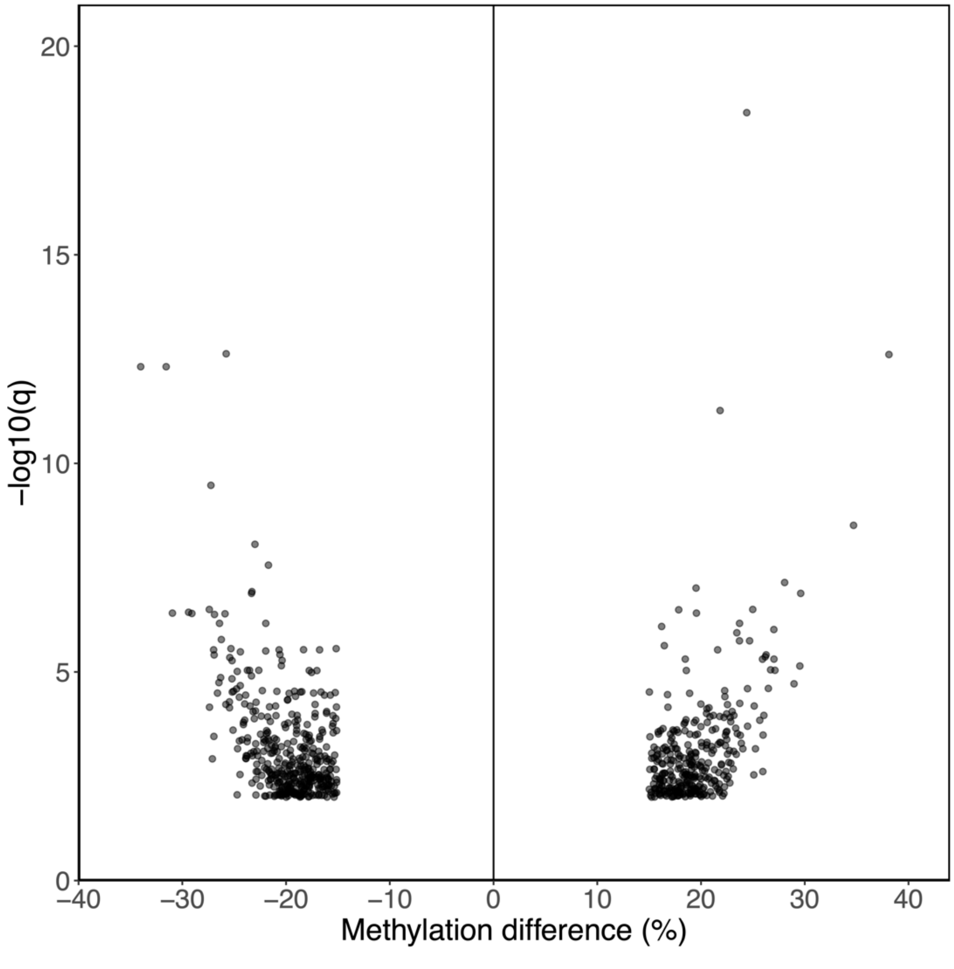

**Supplementary Figure 5.** Manhattan plot of DMS against methylation difference between hatchlings from the different depth treatments. DMS (n=714, methylation difference >15%, q-value < 0.01) are plotted against the methylation difference (%), split between sites that are hyper-methylated in deep-incubated hatchlings (top) and sites that are hyper-methylated in shallow incubated hatchlings (bottom). The 21 DMS of interest are labelled and the q-value is represented by the colour scale.

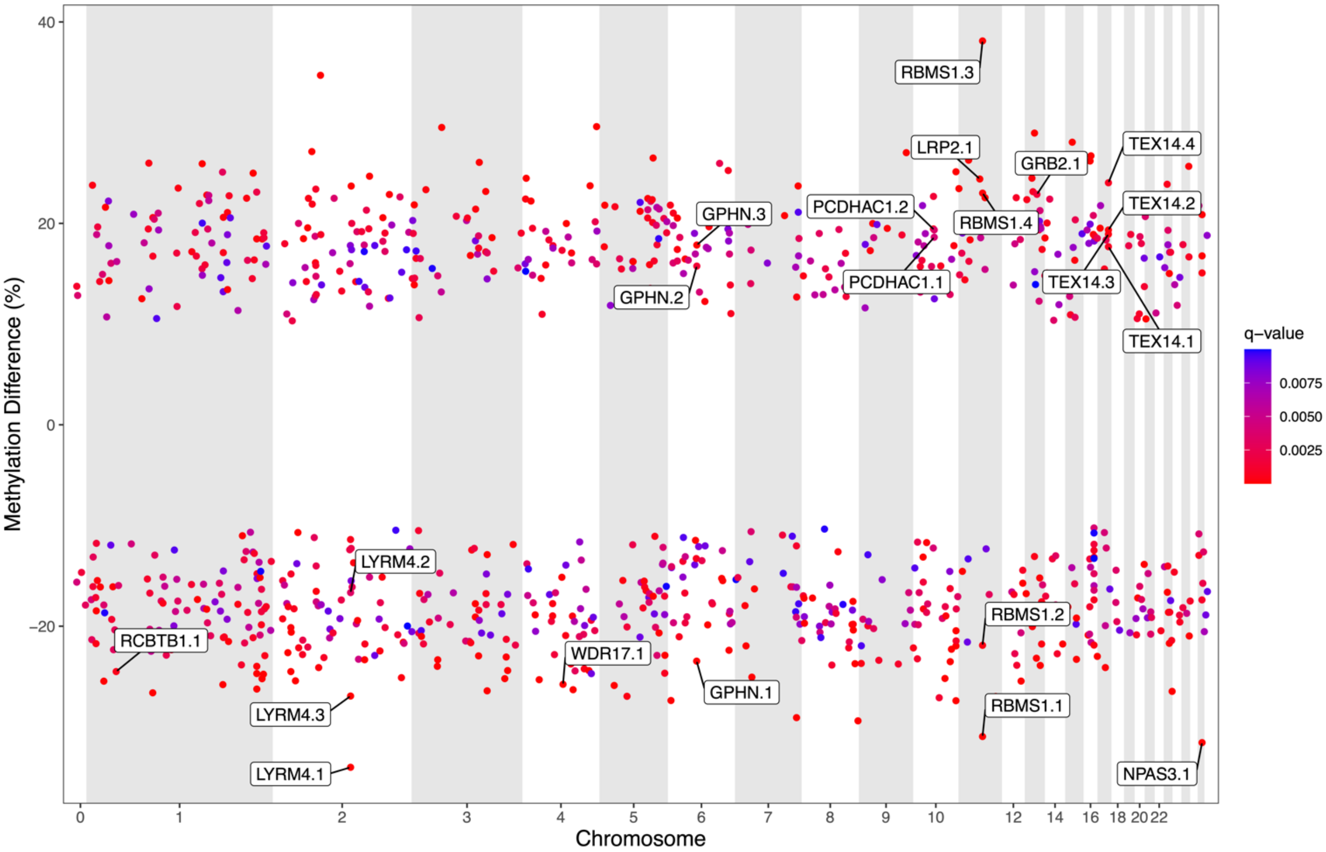

**Supplementary Text 1.** Extended methods for SNP calling pipeline. To obtain a list of C-to-T SNPs to remove from the CpG dataset, the Revelio algorithm was used to mask bases generated by the bisulfite conversion process that may be incorrectly interpreted as SNPs (Nunn et al. 2022), then the GATK pipeline v.4.2.6.1 (McKenna et al. 2010) was used to call SNPs. Read group information was added to the masked, aligned reads with samtools v.1.10 (Li et al. 2010), followed by SNP calling using the GATK pipeline v.4.2.6.1 (McKenna et al. 2010). SNPs were called per sample with HaplotypeCaller. Joint genotyping was performed with GenotypeGVCFs across samples in the deep and shallow treatments separately. This was done to minimise the interference of true differentially methylated sites between depth treatments when calling C-to-T SNPs. This is because siblings in each treatment should have the same genetic background, but have different methylation patterns induced by the incubation condition experienced. Several filtering steps were performed next to give the final SNP dataset. We removed SNPs with (1) a quality by depth less than 2 and mapping quality less than 35 with GATK SelectVariants, (2) coverage in the 99.9$^{th}$ percentile with vcftools v0.1.16 (Danecek et al. 2011), and (3) coverage less than 7 and within 10 bp of an indel with bcftools v1.16 (Danecek et al. 2021). Linkage disequilibrium pruning ($r^2$<0.9 in 10 kb windows) was then applied using bcftools. Finally, SNPs with a minor allele frequency less than 0.05 were removed to produce the final SNP dataset for each treatment.

**Supplementary Table 1.** WGBS statistics per hatchling. For the subset of hatchlings that were selected for sequencing (n=40 total; n=20 per treatment; n=4 per 10 clutches, n=2 per 20 sub-clutches).

| Hatchling ID | Maternal ID | Treatment | Total raw read pairs | Mean mapping efficiency (%) | Bisulfite conversion efficiency (%) | Total CpGs after de-stranding | Mean CpG coverage after de-stranding |
|---|---|---|---|---|---|---|---|
| 162-7 | SLL063 | Shallow | 132125697 | 79.0 | 99.994 | 25420314 | 8.93 |
| 162-11 | SLL063 | Shallow | 138252128 | 80.7 | 99.995 | 24187368 | 7.77 |
| 163-1 | SLL063 | Deep | 132157135 | 74.3 | 99.994 | 25555409 | 9.00 |
| 163-5 | SLL063 | Deep | 138215181 | 80.8 | 99.993 | 23339675 | 7.48 |
| 164-8 | SLL065 | Shallow | 138154759 | 83.3 | 99.995 | 25551989 | 9.75 |
| 164-13 | SLL065 | Shallow | 132236373 | 81.7 | 99.994 | 24235846 | 7.83 |
| 165-1 | SLL065 | Deep | 132286877 | 79.5 | 99.994 | 25522943 | 9.40 |
| 165-3 | SLL065 | Deep | 124309977 | 78.5 | 99.993 | 22879483 | 6.60 |
| 166-12 | SLL146 | Shallow | 132168259 | 78.3 | 99.994 | 25550303 | 9.57 |
| 166-16 | SLL146 | Shallow | 138244303 | 73.6 | 99.993 | 23713831 | 7.16 |
| 167-1 | SLL146 | Deep | 138205348 | 68.2 | 99.992 | 23528252 | 6.68 |
| 167-8 | SLL146 | Deep | 132250210 | 78.2 | 99.994 | 25513139 | 9.06 |
| 168-3 | SLL176 | Shallow | 132310832 | 77.4 | 99.994 | 25489077 | 9.04 |
| 168-4 | SLL176 | Shallow | 139288016 | 82.2 | 99.995 | 25539371 | 10.08 |
| 169-5 | SLL176 | Deep | 138547819 | 80.5 | 99.995 | 25551370 | 9.90 |
| 169-8 | SLL176 | Deep | 132354672 | 75.8 | 99.993 | 25318231 | 8.27 |
| 170-3 | SLL142 | Shallow | 138564032 | 78.8 | 99.995 | 25532667 | 9.76 |
| 170-4 | SLL142 | Shallow | 132251390 | 72.8 | 99.989 | 25434866 | 8.54 |
| 171-1 | SLL142 | Deep | 132277642 | 74.1 | 99.990 | 25426705 | 8.58 |
| 171-2 | SLL142 | Deep | 139003709 | 82.0 | 99.995 | 25511930 | 9.82 |
| 172-1 | SLL188 | Shallow | 132302522 | 75.3 | 99.991 | 25428030 | 8.60 |
| 172-5 | SLL188 | Shallow | 138373670 | 77.1 | 99.992 | 24004894 | 7.42 |
| 173-3 | SLL188 | Deep | 128768337 | 83.4 | 99.994 | 23728787 | 7.37 |
| 173-7 | SLL188 | Deep | 132340278 | 77.9 | 99.992 | 25500965 | 9.10 |
| 174-2 | SLL144 | Deep | 132334347 | 79.7 | 99.991 | 25438012 | 8.91 |
| 174-9 | SLL144 | Deep | 138210202 | 81.8 | 99.994 | 23603725 | 7.58 |
| 175-2 | SLL144 | Shallow | 132255878 | 75.9 | 99.991 | 25487153 | 8.90 |

| | | | | | | | |
|---|---|---|---|---|---|---|---|
| 175-7 | SLL144 | Shallow | 138178365 | 81.4 | 99.995 | 24426063 | 7.71 |
| 176-2 | SLL189 | Shallow | 138698534 | 79.5 | 99.995 | 25541878 | 9.68 |
| 176-5 | SLL189 | Shallow | 132235631 | 78.9 | 99.991 | 25465148 | 8.98 |
| 177-2 | SLL189 | Deep | 128738663 | 75.5 | 99.991 | 25391429 | 8.37 |
| 177-3 | SLL189 | Deep | 138622777 | 82.1 | 99.995 | 25514569 | 9.67 |
| 178-2 | SLL171 | Shallow | 132326833 | 75.3 | 99.991 | 25393011 | 8.48 |
| 178-5 | SLL171 | Shallow | 138221719 | 82.8 | 99.994 | 24782292 | 8.17 |
| 179-1 | SLL171 | Deep | 132307648 | 76.3 | 99.991 | 25488501 | 8.85 |
| 179-6 | SLL171 | Deep | 136149240 | 77.8 | 99.995 | 24299715 | 7.51 |
| 180-6 | SLL143 | Shallow | 132248782 | 78.0 | 99.991 | 25469284 | 8.92 |
| 180-11 | SLL143 | Shallow | 138139674 | 79.9 | 99.995 | 23336240 | 7.33 |
| 181-6 | SLL143 | Deep | 132307312 | 76.7 | 99.992 | 25404232 | 8.58 |
| 181-10 | SLL143 | Deep | 133734003 | 85.1 | 99.996 | 25396507 | 8.96 |
